# Proliferative advantage of specific aneuploid cells drives evolution of tumor karyotypes

**DOI:** 10.1101/2022.04.14.488382

**Authors:** Ivana Ban, Lucija Tomašić, Marianna Trakala, Iva M. Tolić, Nenad Pavin

## Abstract

Most tumors have abnormal karyotypes, which arise from mistakes during mitotic division of healthy euploid cells and evolve through numerous complex mechanisms. In a recent mouse model with high levels of chromosome missegregation, chromosome gains dominate over losses both in pretumor and tumor tissues, whereas tumors are characterized by gains of chromosomes 14 and 15. However, the mechanisms driving clonal selection leading to tumor karyotype evolution remain unclear. Here we show, by introducing a mathematical model based on a concept of a macro-karyotype, that tumor karyotypes can be explained by proliferation-driven evolution of aneuploid cells. In pretumor cells, increased apoptosis and slower proliferation of cells with monosomies lead to predominant chromosome gains over losses. Tumor karyotypes with gain of one chromosome can be explained by karyotype-dependent proliferation, while for those with two chromosomes an interplay with karyotype-dependent apoptosis is an additional possible pathway. Thus, evolution of tumor-specific karyotypes requires proliferative advantage of specific aneuploid karyotypes.

**Significance:** Most tumors have an erroneous number of chromosomes, which arise from mistakes during division of healthy cells and evolve through numerous complex mechanisms, including chromosome missegregation, cell proliferation and cell death. However, understanding the mechanisms leading to tumor evolution from healthy cells is a hot topic. Here we show, by introducing a “macro-karyotype model”, that perturbed number of chromosomes in tumor cells arises predominantly from faster division of cells characterized by a specific combination of chromosomes, or together with irregular cell death. This finding, strengthened by comparison of our theory with experimentally observed combination of chromosomes in different stages of tumor development, gives a direction for future experiments in identifying the key processes underlying tumor development.

## INTRODUCTION

During cell division, missegregation of duplicated chromosomes generates cells with perturbed karyotypes, a condition with abnormal chromosome number known as aneuploidy. Aneuploidy has detrimental effect on cell vitality and proliferation, and is linked with cancer and other diseases (Ben-David and Amon, 2019; Gordon et al., 2012; Hwang et al., 2021; Weaver and Cleveland, 2006). While in healthy cells, aneuploidy causes a strong anti-proliferative response as a result of gene dosage imbalances (Santaguida et al., 2015) this response can be overcome in cancer cells, allowing the increased frequency of errors to promote aneuploid karyotype evolution and acceleration of tumor formation (Duijf and Benezra, 2013; van Jaarsveld and Kops, 2016). These ideas have emerged from extensive studies of aneuploidy in different systems including cell lines (Cimini et al., 2001; Thompson and Compton, 2008, 2011a), organoids (Bolhaqueiro et al., 2019; Drost and Clevers, 2018; Narkar et al., 2021), animal models (Bolton et al., 2016; Sheppard et al., 2012; Shoshani et al., 2021; Trakala et al., 2021), as well as theoretically (Araujo et al., 2013; Desper et al., 2005; Elizalde et al., 2018; Gusev et al., 2000, 2001; Laughney et al., 2015; Lynch et al., 2021). Yet, how the interplay between chromosome missegregation, cell proliferation and other processes drives long-term karyotype evolution is poorly understood.

One of key processes which drives karyotype evolution is chromosome missegregation, which is increased for many tumor karyotypes and termed chromosome instability (CIN) (Hintzen et al., 2021; Nicholson and Cimini, 2013; Thompson and Compton, 2008). Missegregation includes gain or loss of one or a few chromosomes due to incorrect attachment of chromosomes to the mitotic spindle (Bakhoum et al., 2009; Cimini, 2008; Dewhurst et al., 2014; Nicholson et al., 2015a; Thompson and Compton, 2011b). Another key process for karyotype evolution is cell proliferation, which determines how the total number of cells with a certain karyotype changes over time. In particular, it was measured that aneuploid cells typically proliferate slower than euploid cells (Hintzen et al., 2021; Williams et al., 2008). However, certain aneuploid karyotypes with a favorable combination of chromosome gains or losses can lead to enhanced cell proliferation (Ben-David et al., 2014; Heyde et al., 2021; Rohban and Campaner, 2015). In contrast, other aneuploid karyotypes can result in malfunctional cell physiology and consequently apoptosis.

Studying how different processes affect long-term karyotype evolution is experimentally challenging because it requires tracking of all relevant karyotypes over many generations. An alternative approach is mathematical modeling in which simple hypotheses can be explored beyond experimental constraints (Jelenic et al., 2018; Tolic and Pavin, 2021). Modeling is powerful because it allows to identify the dominant mechanisms in the existing experiments and provides ideas for new experiments targeted on testing specific model predictions. Theoretical models of karyotype evolution have been designed to study certain key aspects of this complex process. Early theoretical studies investigated the role of chromosome missegregation on karyotype evolution, leading to a theoretical upper limit for the missegregation rate that allows diploid cells to survive (Gusev et al., 2000, 2001). To study how karyotype evolution depends on genetic linkage, genes controlling cell proliferation, apoptosis and chromosome missegregation were placed on two chromosomes in different combinations (Araujo et al., 2013). A study of karyotype evolution that includes the interplay between missegregation and whole genome duplication found that population converges to near triploid state (Laughney et al., 2015), and that heterogeneity is primary influenced by the missegregation rate (Elizalde et al., 2018). Recently, a theoretical model was developed to measure CIN from karyotype diversity within a tumor (Lynch et al., 2022).

Development of novel mouse models with high levels of chromosome missegregation enabled experimental exploration of karyotype evolution during tumor initiation and growth (Trakala et al., 2021,Shoshani et al., 2021). Quantification of chromosomal gains and losses in these mouse models revealed that chromosome gains dominate over losses both in pretumor and tumor tissues. At early stages of T-cell development in the thymus, 90% of cells are euploid due to strong selection, whereas in later stages cells accumulate more errors and a larger fraction, up to 60-80% of cells become aneuploid, where individual chromosomes are roughly equally gained, on average 4% over aneuploid cells (Trakala et al., 2021). In contrast, when T-cell lymphomas start to develop, there is a dramatic increase in gains of chromosomes 14 and 15 in almost 100% of aneuploid cells, whereas the average gain of other chromosomes remains around 8%. These results suggested that lymphomas developed through clonal selection of favorable aneuploidies. However, the mechanisms of clonal selection leading to tumor karyotype evolution from euploid karyotypes remain unclear.

In this paper we explore the idea that karyotype-dependent proliferation and apoptosis are key drivers of clonal selection that leads to aneuploidy observed in tumor cells. To describe a large number of karyotype combinations that arise due to missegregation, we introduce a “macro-karyotype model” in which we follow the copy number of specific chromosomes, together with the total number of gains and losses for other chromosomes. By solving the model we find that only a small fraction of chromosome missegregation events generates tumor-specific karyotypes, but thanks to their proliferative advantage, cells with these karyotypes eventually overtake the population in agreement with the experimentally observed karyotype evolution. Thus, the central finding of our theory is that, under conditions of high chromosome missegregation, tumor-specific karyotypes arise from proliferative advantage of specific aneuploid karyotypes.

## RESULTS

### Mathematical model for tumor karyotype evolution

Theoretical exploration of tumor karyotype evolution poses two main challenges: tumor tissues consist of a large number of karyotype combinations and a large number of cells. To tackle these problems, we design a novel approach based on a concept of a macro-karyotype. We define a macro-karyotype as a karyotype that contains the copy number of specific chromosomes that are crucial for the process of interest, whereas for other chromosomes only the total chromosome gains and losses are counted. Macro-karyotypes contain all relevant information about the exact cell karyotype, while substantially reducing the number of karyotype combinations and allowing for the use of mean-field calculations, which can describe large populations. Thus, the strength of this approach lies in that it allows us to follow karyotype evolution for a virtually unlimited number of cells and generations.

In our macro-karyotype model we calculate karyotype evolution and a corresponding pedigree tree (**Figure 1A**). Karyotype of a cell with *n* different chromosomes is described by a vector, **K** ≡ (*c*_1_, …, *c*_*n*_ ), in which *c*, denotes copy number of *i*-th chromosome (**Figure 1B**). Because some karyotypes are frequently used, we introduce a notation for them. First, diploid karyotype is denoted as a vector **K**_2*n*_ ≡ (*c*_1_ = 2, …, *c*_*n*_ = 2). Second, karyotypes with at least one chromosome loss form a set of karyotypes *K*_−_ = {(*c*_1_, …, *c*_*n*_ )| ∃ *c*_*i*_ = 1, *i* = 1, …, *n*}, whereas karyotypes with chromosome gains form a set *K*_+_ = {(*c*_1_, …, *c*_*n*_ )| ∃ *c*_*i*_ ≥ 3, *i* = 1, …, *n*}. Finally, we introduce a notation for a set of karyotypes with gain of chromosome 15, *K*_+ 15_ = {(*c*_1_, …, *c*_*n*_)| *c*_15_ ≥ 3}, and a set of karyotypes with gain of chromosomes 14 and 15, *K*_+ 14,+ 15_ = {(*c*_1_, …, *c*_*n*_ )| *c*_14_ ≥ 3, *c*_15_ ≥ 3}. Note that karyotypes with chromosome losses and gains belong to more than one set.

**Figure 1.**
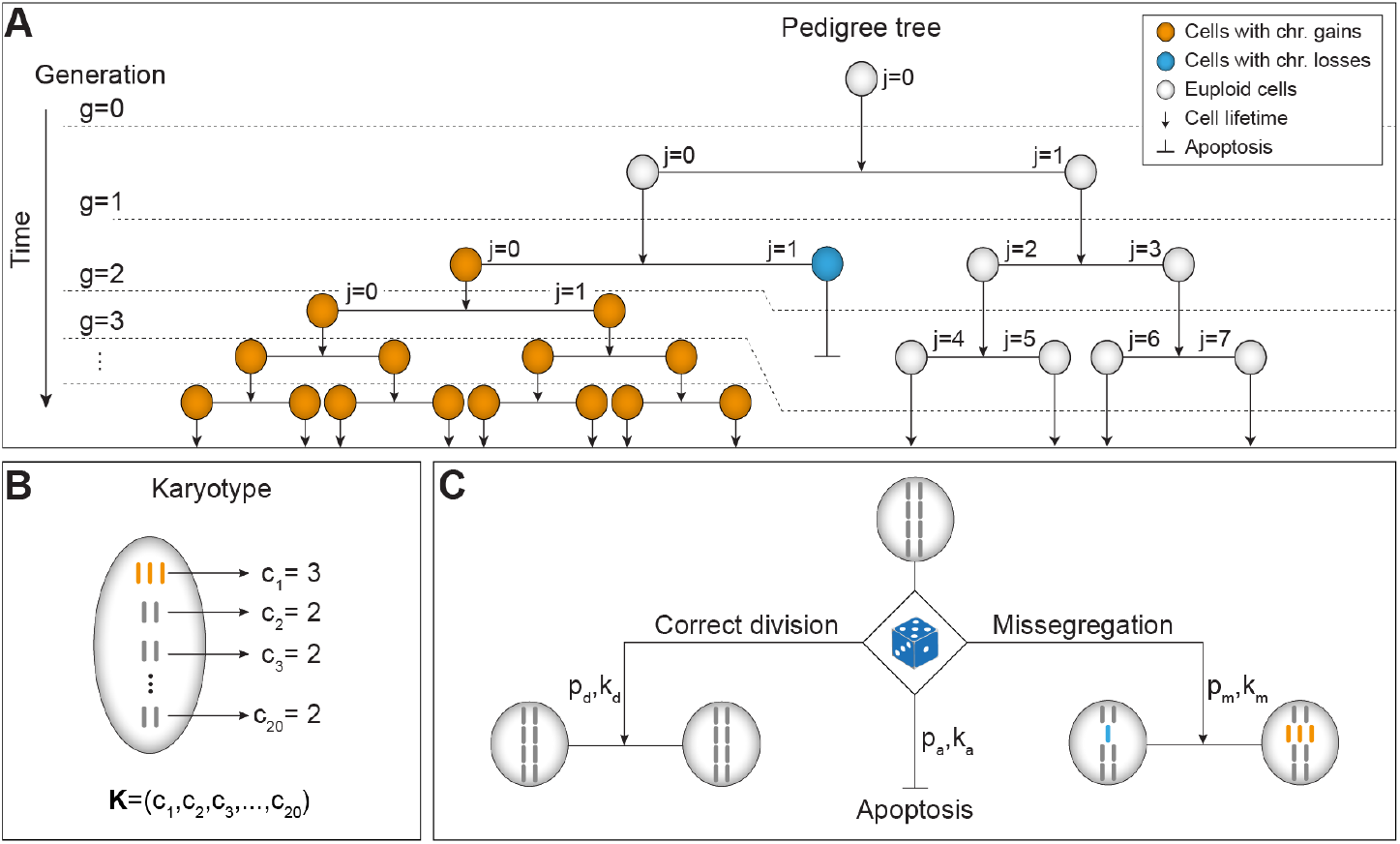
Mathematical model for tumor karyotype evolution. **A**, Scheme of the pedigree tree. Mother cell (gray circle) in generation *g* = 0 and with cell index *j* = 0 divides faithfully to two daughter cells (gray circles) with cell indexes *j* = 0 and *j* = 1 in generation *g* = 1. A chromosome missegregation event occurred yielding one daughter cell with trisomy (orange, *j* = 0) and the other with monosomy (blue, *j* = 1) at generation *g* = 2. Cells with trisomy of specific chromosome divide faster which is represented by a shorter arrow. Cells can also undergo apoptosis (*g* = 2, *j* = 1). Generations are visually divided by horizontal dashed lines which can be cascading due to different proliferation rates for diploid and aneuploid cells. **B**, Mathematical description of cell karyotype. Cell (gray oval) contains *n* chromosomes (rods). First chromosome has 3 copies (orange rods, *c*_1_ = 3), and the other chromosomes have 2 copies (gray rods, *c*_2_ = *c*_3_ = ⋯ = *c*_*n*_ = 2). Karyotype of a cell is fully described by *c*_1_, …, *c*_*n*_. **C**, Processes included in our model. Mother cell probabilistically (dice) undergoes one of 3 processes: correct division, missegregation or apoptosis, with respective probabilities *p*_d_, *p*_m_, *p*_a_ and respective rates *k*_d_, *k*_m_, *k*_a_. Only 4 chromosomes depicted.

By constructing a pedigree tree, we follow multiple generations of cells, starting from a founder cell in generation *g* = 0 (**Figure 1A**). In generation *g* the number of cells (branches) can be up to 2^*g*^, where each cell (branch) has a unique index *j* = 0, …, 2^*g*^ − 1. Because some branches of the pedigree tree are pruned when a cell undergoes apoptosis, cells with corresponding index *j* are missing and the total number of cells in that generation is smaller than 2^*g*^.

Cell lifetime, *t*_0_, depends on the karyotype and ends with either division or apoptosis, which occur with probabilities 1 – *p*_a_ and *p*_a_, respectively (**Figure 1C**). During cell division chromosomes can divide correctly, where each half of the duplicated chromosome (chromatid) goes to one of the two daughter cells, or missegregate, where both chromatids end up in the same cell. These events depend on the karyotype and occur with probabilities *p*_d_ and *p*_m_, respectively (**Figure 1C**). In our model, chromosome missegregation events are independent and the missegregation probability per cell is calculated as 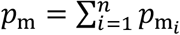. This approximative expression holds for rare events, *p*_m_ ≪ 1. Missegregation probability of the *i*-th chromosome, 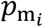, is calculated as the copy number of that chromosome multiplied by its probability to missegregate, 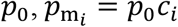. These probabilities, together with cell lifetime and their positions in a pedigree tree, define a discrete stochastic model for karyotype evolution.

### Rate equation for population of cells

While the stochastic model can follow each individual cell, the main disadvantage is that it cannot follow all cells for a large number of generations. To circumvent this issue, we develop a mean-field approach, which allows us to start with a large number of cells and follow them over many generations. Here, we first calculate the probability for finding a cell with a given karyotype at the position within the pedigree tree, *P*_*g,j*_ (**K**), by taking into account that cells in the generation *g* appear by division of the cells in the previous generation, *g* − 1, whereas the cells in the generation *g* disappear upon cell division or apoptosis. To calculate how these probabilities evolve in time, *t*, we introduce the following equation:

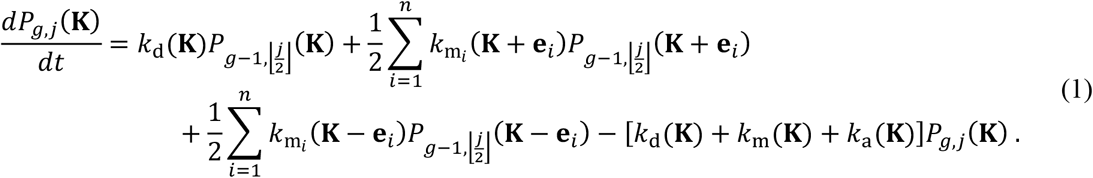

Here, the first term on the right-hand side describes how a cell with karyotype **K** in generation *g* − 1, which we refer to as the mother cell, correctly divides into two cells of the same karyotype in generation *g*, which we refer to as daughter cells. The second and third terms describe missegregation in which the mother cells with karyotypes **K** ± **e**_*i*,_ lose or gain the *i*th chromosome, yielding one daughter cell with karyotype **K**. The last term describes division or apoptosis of the daughter cell and therefore appears with a negative sign. The unit vector **e**, has the value 1 in the *i*th coordinate and 0’s elsewhere.

Indices in Eq. (1) describe the position in the pedigree tree, where each daughter which belongs to generation *g* and branch *j*, originates from a mother cell in generation *g* − 1 with the corresponding branch ⌊*j*/2⌋ (**Figure 1A**). The floor function ⌊ ⌋ ensures that two daughter cells correspond to one mother cell, one daughter cell with even and the other with odd index *j*. In the case of missegregation, each daughter cell has the same probability to acquire one chromosome extra or less. However, only the daughter cell with karyotype **K** will contribute to Eq. (1) and therefore we multiply these terms by 1/2.

The rates in Eq. (1) describe correct cell division, *k*_d_ (**K**), apoptosis, *k*_a_ (**K**), and chromosome missegregation, 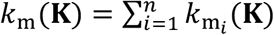, where 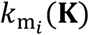 denotes the missegregation rate of the *i*th chromosome (**Figure 1C**). The rates of apoptosis and missegregation of the *i*th chromosome are calculated as 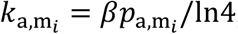, where the proliferation rate relates to the lifetime of a cell as *β* = l*n* 2/*t*_0_. The corrective factor l*n* 4 appears because cell division in Eq. (1) is considered as a Poisson process, where every cell divides with equal probability independent of its age, whereas in more realistic case all cells live equally long (see SI Appendix). The rate of correct division is implicitly given by *β* = *k*_d_ + *k*_m_ + *k*_a_.

To understand karyotype evolution it is important to calculate the number of cells of a certain karyotype in the population over time, irrespective of the position in the pedigree tree. Thus, we define this number as 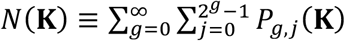 and calculate how it changes in time by summing Eq. (1) over indices *g* and *j* (see SI Appendix),

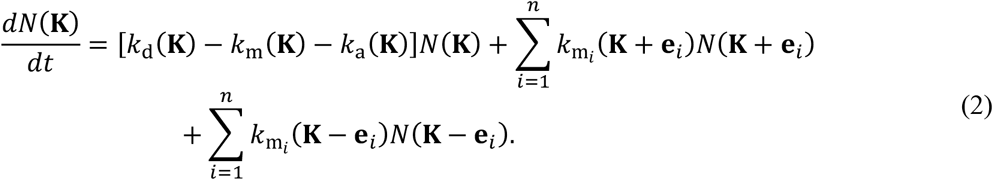

The dimensionality of this rate equation is 2^*g*+1^ − 1 times lower than that of Eq. (1), yet it retains the complete information about the karyotypes in a cell population.

Despite the reduced dimensionality, Eq. (2) cannot be directly integrated on a supercomputer because of a huge number of karyotype combinations. For this reason, we solve Eq. (2) by reducing the space of vector **K** while retaining all the important information about the karyotype, e.g., whether it is euploid or aneuploid, whether aneuploid karyotypes contain monosomies and trisomies. Thus, we construct a vector that contains the information about the karyotype, **M**(**K**) ≡ (*x*_1_, …, *x*_*L*_), which we term macro-karyotype. Here, component 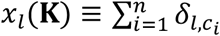 denotes the number of chromosomes with *l* copies, and index *L* denotes the maximal copy number of each chromosome. Because nullisomies are not described in our model, the number of different chromosomes per cell is conserved, *x*_1_ + ⋯ + *x*_*L*_ = *n*. Thus, the macro-karyotype defined in this manner includes multiple karyotypes. A certain macro-karyotype is obtained as a set of permutations of vector components for each chromosome *c*_*i*_ and the number of cells with this macro-karyotype is *Ñ* (**M**) = (*x*_1_ ! ⋯ *x*_*L*_ !)^−1^ × ∑_all perm._ *N*(**K**). Note that the normalization is correction for counting the same karyotype multiple times.

In order to calculate how the number of cells with a certain macro-karyotype changes in time, we sum Eq. (2) over all permutations reaching the final form of the rate equation for macro-karyotype evolution (see SI Appendix):

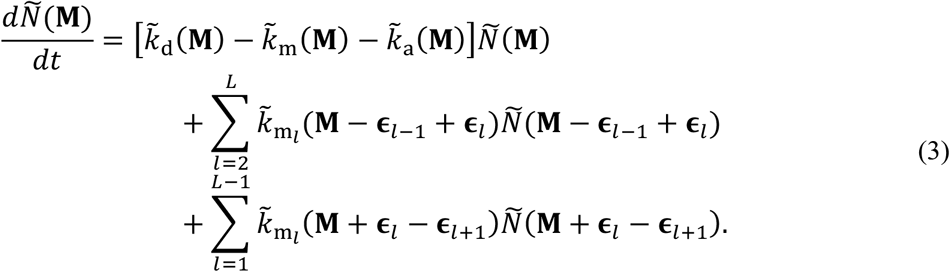

This rate equation is derived with the assumption that rates *k*_d,m,a_ (**K**) have the same value for all permutations of *c*_1_, …, *c*_*n*_ that belong to the same macro-karyotype **M**, and thus the rates obey 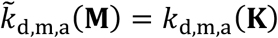, except in the case where 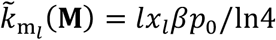. The unit vectors ε_*l*_, ε_*l* + 1_ and ε_*l* − 1_ have the value 1 in the *l*th, *l* + 1 and *l* − 1 coordinate, respectively, and 0’s elsewhere. For studying karyotypes with a specific chromosome, we generalize our approach by constructing an extended macro-karyotype **M**_ext_ (**K**) ≡ (*x*_1_, …, *x*_*L*_, *c*_15_), where the last component, which corresponds to the specific chromosome, accounts for number of copies of chromosome 15 (see SI Appendix).

The key advantage of the macro-karyotype approach is the substantially reduced dimensionality of the vector space, in comparison with the dimensionality of the original vector space. For example, in the case of a cell with *L* = 6 and *n* = 20, the number of possible macro-karyotypes is *L*(*n* − 1)^*L* − 1^ ≈ 1.5 · 10^7^, which is 8 orders of magnitude smaller than the number of possible karyotypes, *L*^*n*^ ≈ 3.7 · 10^15^. Thus, Eq. (3) is a unique mathematical tool that allows us to study tumor karyotype evolution because it describes a large number of cells and karyotype combinations.

### Increased apoptosis and slower proliferation of cells with monosomies lead to predominant chromosome gains over losses

In order to understand the role of apoptosis and proliferation in tumor karyotype evolution, we solve our model and compare the obtained results with the experimentally found karyotypes in pretumor and tumor cells from a mouse model (Trakala et al., 2021). First, we explore under what conditions chromosome gains dominate over losses in pretumor cells, and subsequently how tumor-specific karyotypes arise.

To explore macro-karyotype evolution starting from multiple diploid cells (**Figure 2A**), we solve Eq. (3), for which we first have to estimate model parameters (see Table 1). For euploid cells, we use the proliferation rate value *β*(**K**_2*n*_ ) = l*n* 2 d^−1^, based on division time for the earliest thymocyte, double negative, stages (DN1-DN4) (Petrie and Zúñiga-Pflücker, 2007). For aneuploid cells, the proliferation rate is *β*(**K** ≠ **K**_.2*n*_ ) = 0.8 l*n* 2 d^−1^, because aneuploid cells proliferate roughly 20% slower in comparison to diploid cells (Hintzen et al., 2021; Williams et al., 2008). Given that in this mouse model (Trakala et al., 2021) all cells have the same probability of missegregating their chromosomes independent of their karyotype, we use the same missegregation rates values for all karyotypes, including both euploid and aneuploid cells. Because there is no direct measurement of the missegregation rate for this mouse model, we roughly estimate it from Eq. (2) in the regime of small chromosome gains, *g*a*in* ≪ 100%, and small number of generations in which linear relationship holds *g*a*in* = *p*_0_ *t*/*t*_0_ (see SI Appendix). Therefore, the missegregation rate is obtained from linear fit of experimentally measured chromosome gains through DN1-DN4 stages, yielding *p*_0_ = 0.0017 (**Figure 2B, Table 1**). Since the rate of apoptosis is not measured for this mouse model, we use it as a free parameter and estimate it later by comparing results of our model with experiments.

**Table 1.**
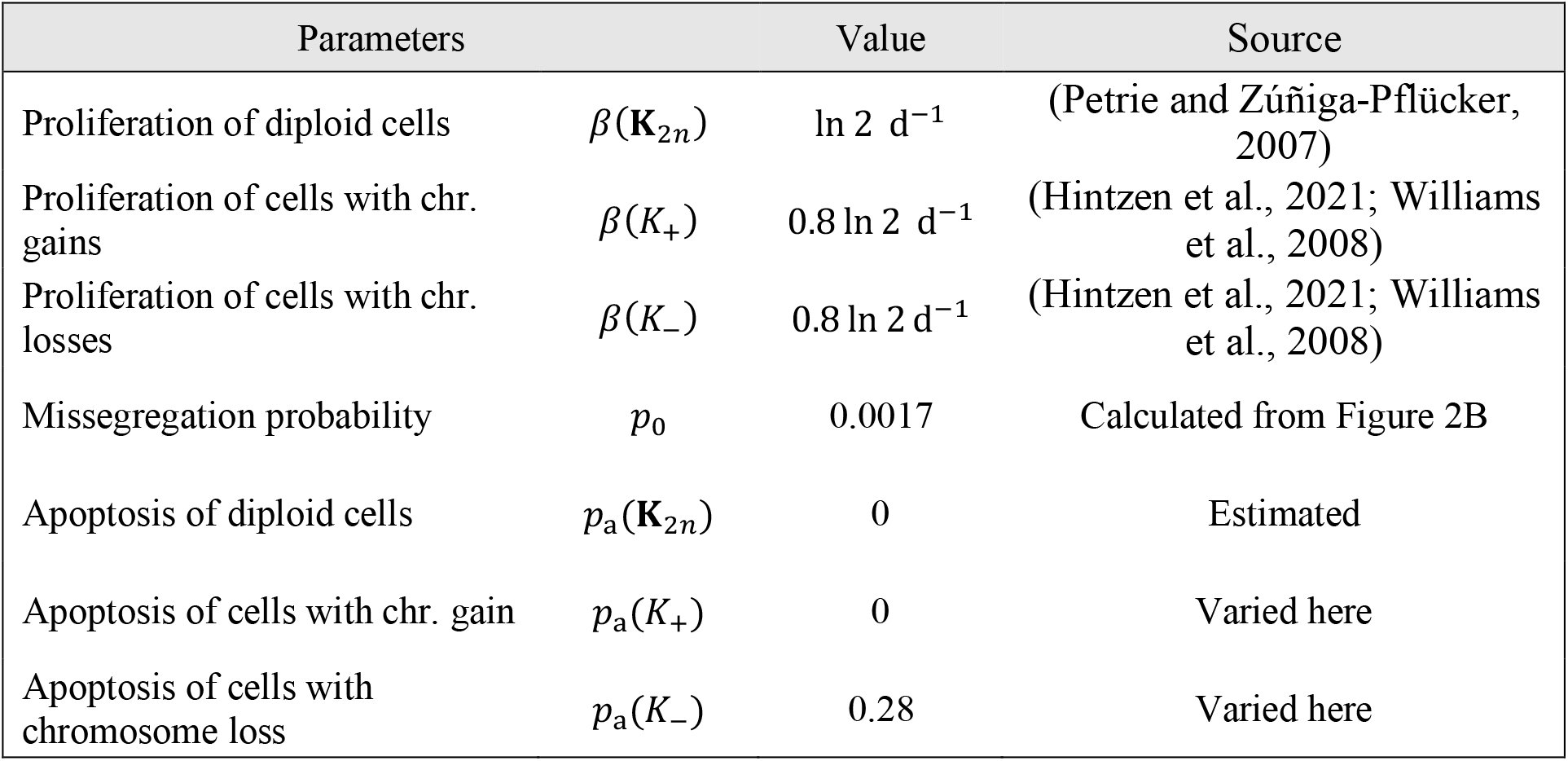
Parameters used in the model.

**Figure 2.**
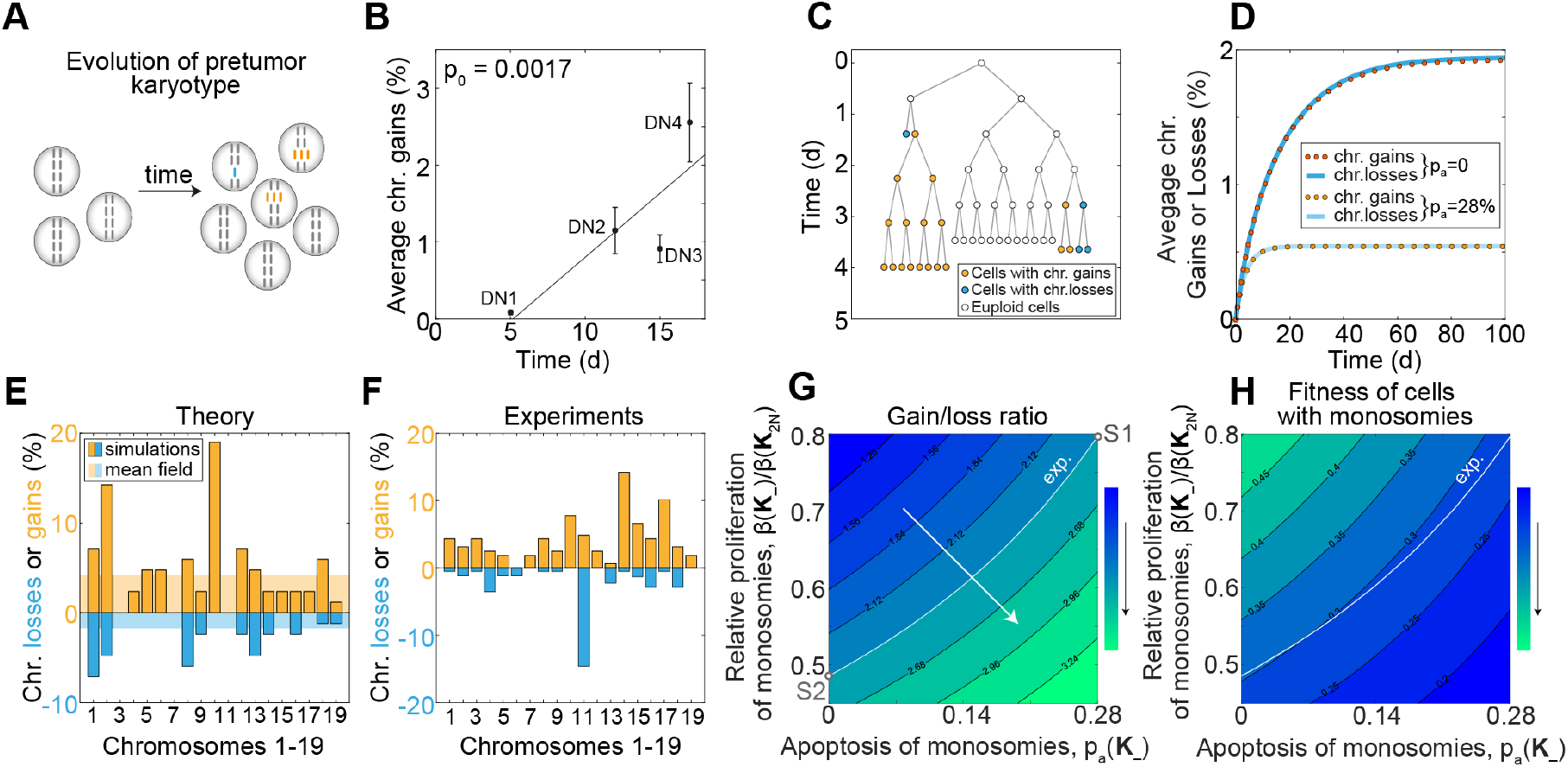
Increased apoptosis and decreased proliferation of cells with monosomies generate imbalance of chromosome gains and losses. **A**, Schematics showing cells containing one copy of chromosome (blue rod), two copies of chromosome (gray rods) and three copies of chromosome (orange rods) at two different time points. **B**, Experimentally measured average chromosome gains during T-cell lymphoma development, for stages DN1-DN4 (black circles) with duration 10 days, 4 days, 2 days, and 2 days, respectively (Porritt et al., 2003). The experimental points are positioned at the middle of duration of each phase. The line represents linear fit, gain = *p*_0_ *t* + offset. **C**, Pedigree tree obtained from one simulation run, showing diploid cells (white circles), cells with gains (orange circles) and losses (blue circles). **D**, Average chromosome gains (dotted orange lines) and losses (full blue lines) in time calculated by mean field approach for *p*_a_ (**K**) = 0 (dark orange and dark blue) and *p*_a_ (**K** ≠ **K**_2*n*_ ) = 28% (light orange and light blue). **E**, Percentage of chromosome gains (orange) and losses (blue) calculated by stochastic simulations (bars, gains 4.7 ± 4.9%, losses 1.8 ± 2.3 %) and mean field model (transparent area, gains 4.2%, losses 1.8%) after *t* = 17 d. Results are mean ± standard deviation if not stated otherwise. Chromosome gains and losses are normalized by number of aneuploid cells. **F**, Percentage of chromosome gains (orange bars, 4.3 ± 3.4%) and losses (blue bars, 1.8 ± 3.3%) experimentally measured in DN4 stage of aneuploid thymus cells (Trakala et al., 2021). **G** and **H**, Chromosome gain/loss ratio and cell fitness (color plots, values denoted along black lines) for different values of proliferation and apoptosis of monosomies. Black arrow besides the color bar denotes increasing trend. In panel **G** transition from small to high gain/loss ratio (from blue to green) is denoted by the white arrow, experimentally measured gain/loss ratio is 2.4 (white line). Parameters corresponding to scenario S1 and S2 are depicted (white circles). In panel **H** white line represents experimentally measured gain/loss ratio from panel **G**. Parameters in panels **C, D, E, G, H** are given in **Table 1** unless stated otherwise.

To visualize karyotype evolution, we plot a pedigree tree for the first 5 generations obtained from one run of stochastic simulations starting with one diploid cell (**Figure 2C**). After several generations missegregation events occur yielding daughter cells with chromosome loss and chromosome gain. Aneuploid cells need more time to divide and therefore their branches are longer in comparison to branches of euploid cells. Some cells undergo apoptosis which is visible as pruned branches in the pedigree tree.

To explore how karyotypes evolve, we solve Eq. (3) and plot chromosome gains and losses in time. We find that chromosome gains and losses closely follow each other and reach a similar asymptotic value (**Figure 2D**). In contrast, experimental data show two times more cells with chromosome gains in comparison to cells with chromosome losses (Trakala et al., 2021). To explain these experimental results, we explore two scenarios: cells with monosomies undergo increased apoptosis (S1) or cells with monosomies proliferate slower (S2), in comparison to cells with chromosome gains. To test scenario S1, we use the probability of apoptosis for monosomies *p*_a_ (**K**_−_) = 28%, whereas all other cells do not undergo apoptosis. We explore the karyotypes after *t* = 17 d, because this time point corresponds to stage DN4, the last stage with no significant gain of specific chromosomes. The model predicts on average 4.2% of cells with chromosome gains and 1.8% of cells with chromosome losses (**Figure 2E**). Similar results are obtained by analyzing karyotypes of 125 cells randomly chosen from 5 runs of stochastic simulations, which also yield the information about the variability in chromosome gains and losses (**Figure 2E**).

By comparing our theoretical results with the experimentally measured gains and losses for thymus cells in DN4 stage (**Figure 2F**) we find that the model quantitatively explains the both average chromosome gains and losses as well as the variability of these distributions (two sample Kolmogorov-Smirnov test, *p*-value is 0.5 for chromosome gains and losses). Because experimentally observed gains and losses can be explained with the model with increased apoptosis of cells with monosomies, we conclude that gain/loss ratio can arise because of karyotype dependent apoptosis, supporting scenario S1.

To explore scenario S2, we vary the proliferation rate for vanishing apoptosis (**Figure 2G**, ordinate). As the proliferation rate of cells with monosomies decreases, gain/loss ratio increases and for proliferation of 0.48 d^−1^ the model yields the experimentally observed gain/loss ratio of 2.4 (**Figure 2G**, point S2). However, it is more likely that cells with monosomies have altered both apoptosis and proliferation. Thus, we vary both parameters simultaneously, to explore the interplay between scenarios S1 and S2. We find a smooth transition from small to large gain/loss ratio as apoptosis increases and proliferation decreases (**Figure 2G**, arrow). The experimentally observed gain/loss ratio is obtained for combinations of apoptosis and proliferation rates where both rates either increase or decrease, to compensate the effect of each other (**Figure 2G**, white contour). We expect that increased apoptosis and slower proliferation of cells with monosomies will reduce their fitness, which is generally considered as ability of a cell to reproduce and defined as *W*(**K**) = *N*(**K**)^−1^ *dN*(**K**)/*dt*. We find cells with high fitness in the region of parameter space where chromosome gains are as frequent as losses, and cells with low fitness in the region where gains dominate over losses (compare **Figures 2G** and **2H**). These results lead us to conclude that both increased apoptosis and slower proliferation of cells with monosomies can explain experimentally measured gain/loss ratio, and that they are interchangeable.

### Karyotype-dependent proliferation drives the evolution of a tumor-specific karyotype with gain of chromosome 15

In contrast to pretumor karyotypes, in which there is no bias for gain of any specific chromosome, karyotypes of T-cell lymphomas are characterized by high occurrence of a chromosomes 14 and 15 (Shoshani et al., 2021; Trakala et al., 2021). As the first step in understanding the evolution of this tumor-specific karyotype, we use our model to explore a simple case where a single chromosome, referred to as chromosome 15, appears with substantially larger gain in comparison to other chromosomes, whereas in the second step we explore a more realistic case of simultaneous gains of two chromosomes. We ask under what conditions the model reproduces experimental data for gains of a specific chromosome and all other chromosomes throughout pretumor and tumor stages (**Figure 3A**).

**Figure 3.**
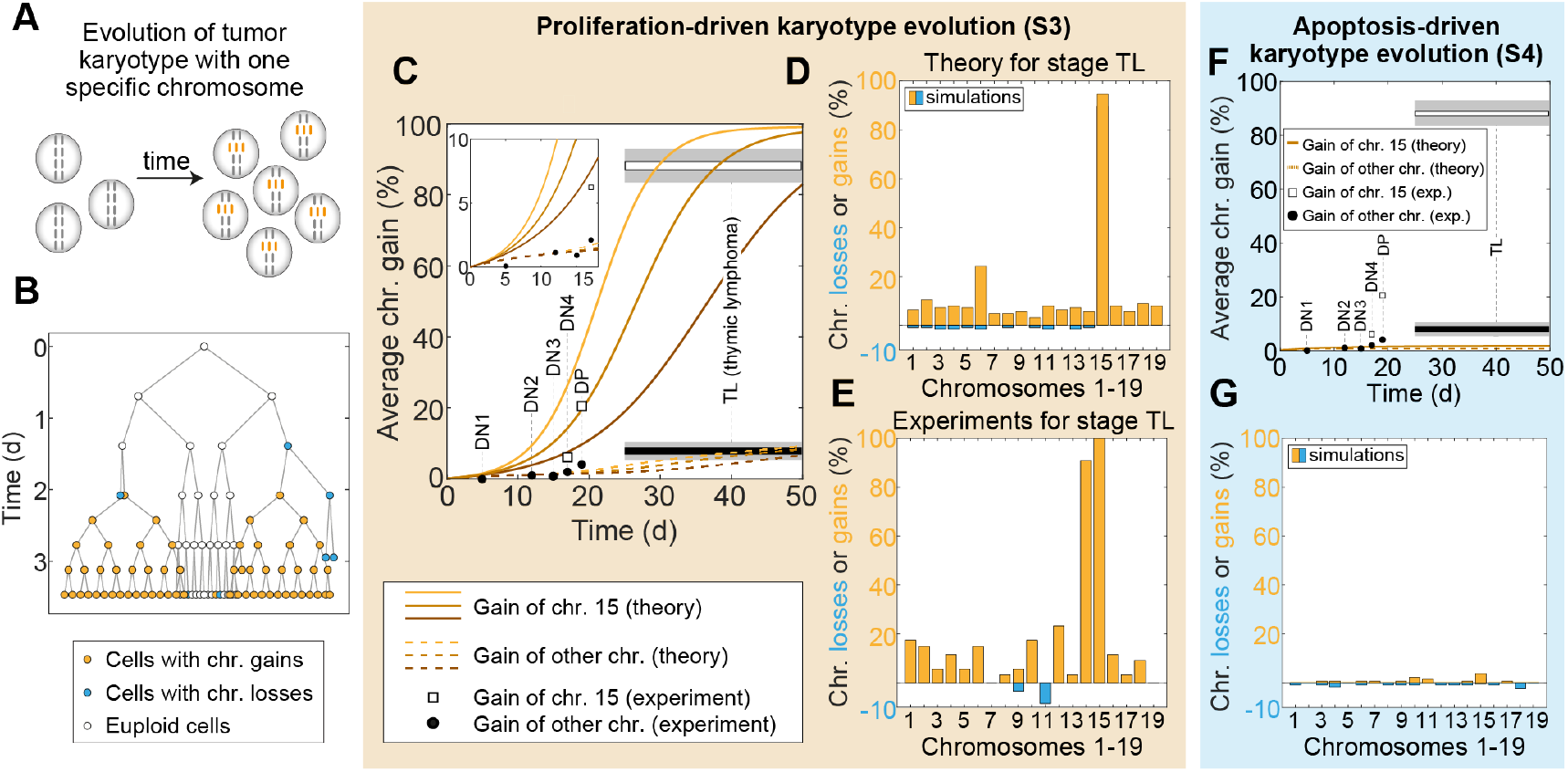
Increased proliferation of cells with trisomy of chromosome 15 drives the evolution of a tumor-specific karyotype. **A**, Schematics showing cells containing two copies of chromosomes (gray rods) and three copies of a specific chromosome (orange rods) at two different time points. **B**, Pedigree tree obtained from one simulation run, showing diploid cells (white circles), cells with gains (orange circles) and losses (blue circles). **C**, Solutions of the mean field model showing time evolution of gain of chromosome 15 (full lines) and gain of other chromosomes (dashed lines) for 3 different proliferation rates, *β*(*K*_+ 15_) = 1.15 d^−1^(dark orange lines), *β*(*K*_+ 15_) = 1.25 d^−1^(orange lines), *β*(*K*_+ 15_) = 1.35 d^−1^(light orange lines). Experimentally measured gains (black circles) in different stages of T-cell lymphomas development. In stages DN4, DP and TL gain of chromosome 15 is depicted separately (white rectangles). Due to experimental uncertainty, values for TL stage are shown for a broad time interval, depicted as long white (chromosome 15), black (other chromosomes) and gray rectangles (standard error of mean). Insert enlarges the first 18 days of T-cell development (DN1-DN4 stages). **D**, Percentage of chromosome gains (orange bars) and losses (blue bars) calculated by stochastic simulations obtained after *t* = 40 d. Parameters same as for middle orange line in panel **C**. Gain of chromosome 15 is 94.4% and loss is 0%, whereas average gain of other chromosomes is 7.8 ± 4.4% and average loss is 0.72 ± 0.67% **E**, Percentage of chromosome gains (orange bars) and losses (blue bars) experimentally measured in T-cell lymphomas (Trakala et al., 2021). Average gain of chromosomes 14 and 15 is 95.4 ± 6.5% and average loss is 0%, whereas average gain of other chromosomes is 8.7 ± 7.1% and average loss is 0.7 ± 2.1%. **F**, Solutions of the mean field model showing time evolution of gain of chromosome 15 (full line) and gain of other chromosomes (dashed line) for parameters *p*_*a*_ (*K*_+ 15_) = 0 and *p*_a_ (**K** ≠ **K**_2*n*_ ) = 0.28. Experimental points are same as in panel **C. G**, Percentage of chromosome gains (orange bars) and losses (blue bars) calculated by stochastic simulations for parameters from panel **F**. Parameters in panels **B, C, D, F** and **G** are given in **Table 1** unless stated otherwise.

In a manner analogous to theory for pretumor karyotype evolution, we explore 2 scenarios: cells with gain of chromosome 15 have increased proliferation (S3) or decreased apoptosis (S4) in comparison to other aneuploid karyotypes. To test scenario S3, we varied the proliferation rate of these cells. A pedigree tree depicts how faster proliferation leads to domination of cells with gain of chromosome 15 (**Figure 3B**). The whole evolution of this macro-karyotype over pretumor and tumor stages is obtained by solving Eq. (3) (**Figure 3C**). For the first 2-3 days, gain of chromosome 15 and gain of other chromosomes follow a similar trend. Later in time, gain of chromosome 15 increases faster than gains of other chromosomes. Strikingly, after 20 days, 10% – 40% cells gained chromosome 15 and after 40 days this fraction reaches 60% – 98%. In contrast, gain of other chromosomes is below 10% for all time points.

These theoretical results are in agreement with experiments, which showed that gain of chromosomes 15 is around 90% whereas gain of other chromosomes is around 10% (**Figure 3C**). This pronounced gain of specific chromosomes is a robust result of our theory because this gain appears for different proliferations and persists afterwards. Our theory also reproduces the main features of karyotype evolution during earlier stages of T-cell development, including steady increase in chromosome gains during DN1-DN4 stages followed by a sharper increase in double positive (DP) stage (**Figure 3C**), supporting the idea of proliferation-driven karyotype evolution.

We also tested robustness of our mean-field result by performing stochastic simulations that corresponds to a realistic situation with a finite number of cells. Indeed, stochastic simulations after 40 days showed same trends as mean-field theory, with pronounced gain of a specific chromosome and lower average gain of other chromosomes (compare **Figures 3C** and **3D**). In experiments it was found that T-cell lymphomas were highly aneuploid and almost all cells contained gain of chromosome 15 (**Figure 3E**). By comparing the distributions obtained by stochastic simulations and experiments, we find that they are different for chromosome losses, but not for chromosome gains (two sample Kolmogorov-Smirnov test, *p*-value is 0.2 for chromosome gains and 0.03 for chromosome losses). The agreement between experiments and theory suggests that increased proliferation of cells with gain of chromosome 15 is the main reason for gain of specific chromosomes in tumor karyotypes.

To test scenario S4 in which cells with gain of chromosome 15 have decreased apoptosis, we set that these cells and euploid cells do not undergo apoptosis, whereas the probability of apoptosis is 28% for other aneuploid cells. In contrast to scenario S3, which resulted in a pronounced gain of chromosome 15 (**Figure 3C**), we find that in scenario S4 all gains stay at a low value, below 2% (**Figure 3F**). Thus, the karyotype distribution obtained by these parameters cannot reproduce the experimentally observed high occurrence of chromosome 15 (**Figure 3G**). In particular, although this theoretical result can explain early stages of T-cell evolution, there is a big discrepancy with experiments for later stages. Therefore, we conclude that reduced apoptosis of cells with gain of chromosome 15 cannot explain the experimentally observed karyotype evolution, additionally strengthening the idea that this process is driven by increased proliferation of these cells.

### Tumor karyotypes with gains of two chromosomes arise due to karyotype-dependent proliferation alone or together with karyotype-dependent apoptosis

To explore the case of simultaneous gains of two chromosomes (**Figure 4A**), as observed in experiments, we test the scenario where cells with simultaneous gains of both specific chromosomes, 14 and 15, have proliferative advantage compared to all other karyotypes (S5) or scenarios in which these two chromosomes regulate different processes, apoptosis (S6) or missegregation rate (S7), in combination with proliferation. In scenario S5, where the proliferation rate of cells with gains of chromosomes 14 and 15 is increased (**Figure 4B, 4C**), we find that during the first 10 days all chromosomes follow a similar trend, whereas later in time gains of chromosomes 14 and 15 increase fast and approach 100%, while the gains of other chromosomes remain at around 10%. The general trend in **Figure 4B** is similar to the scenario S3 based on increased proliferation for gains of a single chromosome (**Figure 3C**), but in **Figure 4B** we observe that gain of specific chromosomes occurs later in time in comparison to **Figure 3C**. This time lag occurs because it is less likely to gain two specific chromosomes than only one. Interestingly, even though we required that karyotypes have gain of two specific chromosomes, which seems as a more rigid constraint than gain of only one chromosome, the agreement between experiments and theory additionally improves (compare **Figures 3C** and **4B**). These results suggest that enhanced proliferation of cells with gain of both specific chromosomes can drive development of T-cell lymphomas.

**Figure 4.**
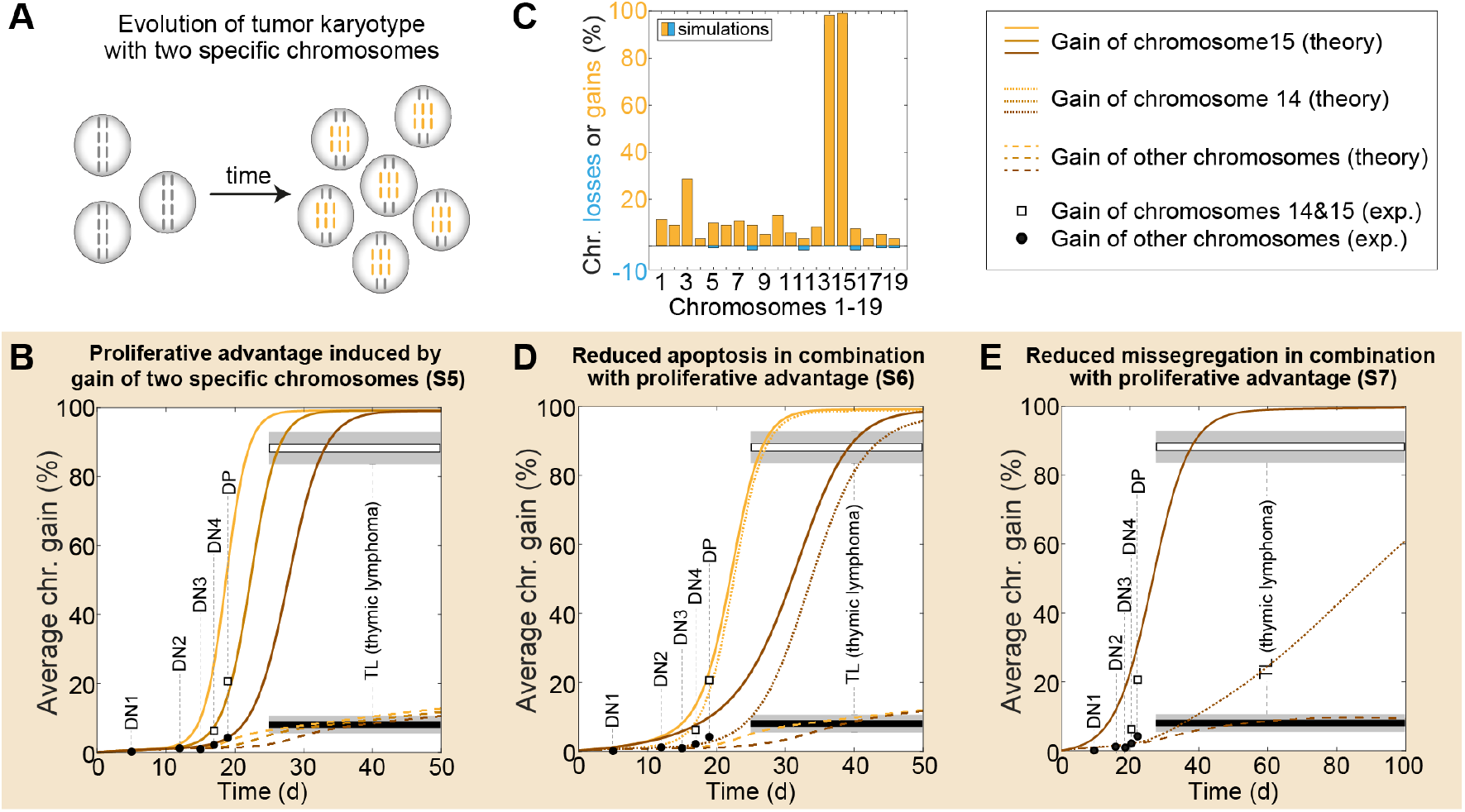
Tumor karyotypes with gains of chromosomes 14 and 15 arise due to karyotype-dependent proliferation alone or together with karyotype-dependent apoptosis. **A**, Schematics showing cells containing two copies of chromosomes (gray rods) and three copies of two specific chromosomes (orange rods) at two different time points. **B**, Solutions of the mean field model showing time evolution of gain of chromosomes 14 and 15 (full lines) and gain of other chromosome (dashed lines) for 3 different proliferation rates, *β*(*K*_+14,+15_) = 1.6 d^−1^ (dark orange lines), *β*(*K*_+14,+15_) = 1.8 d^−1^ (orange lines), *β*(*K*_+14,+15_) = 2.0 d^−1^ (light orange lines). **C**, Percentage of chromosome gains (orange bars) and losses (blue bars) calculated by stochastic simulations after *t* = 40 d for parameters from panel **B** (middle lines). **D** and **E**, Solutions of the mean field model showing time evolution of gains of chromosome 14 (dotted lines), chromosome 15 (full lines) and other chromosomes (dashed lines). In panel **D**, two different probabilities of apoptosis *p*_a_ (*K*_+14_) = 0.028 (orange lines) and *p*_a_ (*K*_+14_) = 0.14 (dark orange lines). Apoptosis of other aneuploid karyotypes is set to *p*_a_ (**K** ≠**K**_2*n*_) = 0.28 and proliferation rate is *β*(*K*_+15_) = 1.8 d^−1^. In panel **E**, missegregation probability for cells with gain of chromosome 14 is *p*_0_ (*K*_+14_) = 0.0001 and the proliferation rate is *β*(*K*_+15_) = 1.25 d^−1^. Experimental points in panels **B, D** and **E** are same as in **Figure 3C**. Parameters in panels **B**-**E** are given in **Table 1** unless stated otherwise.

In scenario S6, we explore the case in which cells with gain of chromosome 14 have decreased apoptosis, whereas cells with gain of chromosome 15 have increased proliferation. As expected, increased proliferation of chromosome 15 leads to its fast appearance (**Figure 4D**). Surprisingly, we find that chromosome 14 also appears with high gains following closely gain of chromosome 15, which is dramatically different from results obtained in S4 where decreased apoptosis alone was not enough to reproduce experimental points at thymic lymphoma. Theoretically predicted high gains of chromosomes 14 and 15 in S6 are consequence of cooperation between increased proliferation and decreased apoptosis in cells with gain of both chromosomes. We also explore whether a combination of missegregation and cell proliferation can explain gains of specific chromosomes (S7). Indeed, decreased missegregation rate of cells with gain of chromosome 14 increases their occurrence in time (**Figure 4E**). However, the increase of chromosome 14 appears later in time compared to chromosome 15 and it is not in agreement with the experimental points at DP stage. Thus, we conclude that simultaneous gain of two specific chromosomes can also appear as a result of increased proliferation in combination with decreased apoptosis.

Taken together, our model predicts that tumor karyotypes characterized by high occurrence of one specific chromosome can be explained by proliferation-driven evolution of aneuploid cells. In the case of tumor karyotypes with two specific chromosomes, the model predicts that such karyotypes can arise by evolution that is driven by karyotype-dependent proliferation or by an interplay of proliferation and apoptosis.

## DISCUSSION

In this study we developed a theoretical model by which we explored karyotype evolution in mouse model with enhanced chromosome missegregation. By quantitative comparison of our theoretical results and experimentally observed karyotypes in different stages of development of thymic lymphomas, we found that cell proliferation plays a key role in selection of a specific chromosome. We also found that in the case with more than one specific chromosome, decreased apoptosis and missegregation can have a role in selection of the specific karyotypes. Because this theory cannot directly identify the key process behind selection of two specific chromosomes, quantitative measurements of parameters for aneuploid karyotypes are necessary (Hintzen et al., 2021). Such measurements in combination with our theoretical predictions will greatly help in understanding tumor karyotype evolution.

Interestingly, we found that specific karyotypes characterized by a decreased missegregation result with increased number of cells with such karyotypes. At first glance, this result contradicts with experimentally observed increased missegregation of various aneuploid karyotypes (Ganem et al., 2009; Lengauer et al., 1997; Thompson and Compton, 2008). Although missegregation of aneuploid karyotype is generally increased, different aneuploid karyotypes are characterized by various missegregation rates (Hintzen et al., 2021; Nicholson et al., 2015b). Our theoretical result, in combination with these experimental findings, suggests that those cells with lower missegregation will be selected among different aneuploid karyotypes.

Numerous studies show that tetraploidization is an important pathway to aneuploidy in human cancer cells, which is based on direct observations of this process in tumor cell lines (Cimini et al., 1999; Shi and King, 2005) and deduced from distribution of chromosome numbers in different tumors (Prasad et al., 2022; Storchova and Kuffer, 2008). In order to investigate theoretically the role of this pathway and its interplay with other processes, our model can be extended by including whole genome duplication as an additional choice of cell fate at each division. Whole genome duplication has been suggested as a key process leading to near triploid karyotypes often observed in tumors (Elizalde et al., 2018; Laughney et al., 2015), but it will be interesting to explore the importance of karyotype dependent proliferation in evolution of these karyotypes.

Karyotypes in human tumors are characterized by the pronounced gains and losses of specific chromosomes, similar to the karyotype patterns studied here (Bolhaqueiro et al., 2019; Liu et al., 2018). Because of the similarities among these karyotype patterns, we expect that the driving mechanism of tumor karyotype evolution identified in this work is directly relevant for human tumors as well. To explore this exciting idea, it will be crucial to implement “macro-karyotype model” to human tumors, which will give a direction for future experiments in identifying the key processes underlying tumor development.

## AUTHOR CONTRIBUTIONS

N.P. designed and supervised this study. I.B., L.T. and N.P. developed the theory. I.T. and M.T. contributed with ideas and discussions. I.B. and N.P. wrote the manuscript with input from L.T. and I.T.

## ACKNOWLEDGEMENTS

We thank Kruno Vukušić for valuable discussions on tumor karyotype evolution and help in comparing experiments and theory; Veronika Pisačić for comments on the manuscript; Iva Dundović, Patrik Risteski, Maja Novak and the rest of the N.P and I.T. groups for discussion and advice; and Ivana Šarić for editing the figures. Research is supported by the European Research Council (Synergy Grant, GA number 855158, granted to I.M.T. and N.P.) and the QuantiXLie Center of Excellence, a project co-financed by the Croatian Government and European Union through the European Regional Development Fund—the Competitiveness and Cohesion Operational Programme (grant KK.01.1.1.01.0004).

## SUPPLEMENTAL INFORMATION

### Karyotype rate equation

In order to calculate how number of cells with certain karyotype changes in time we sum Eq. (1) over indices *g* and *j*,

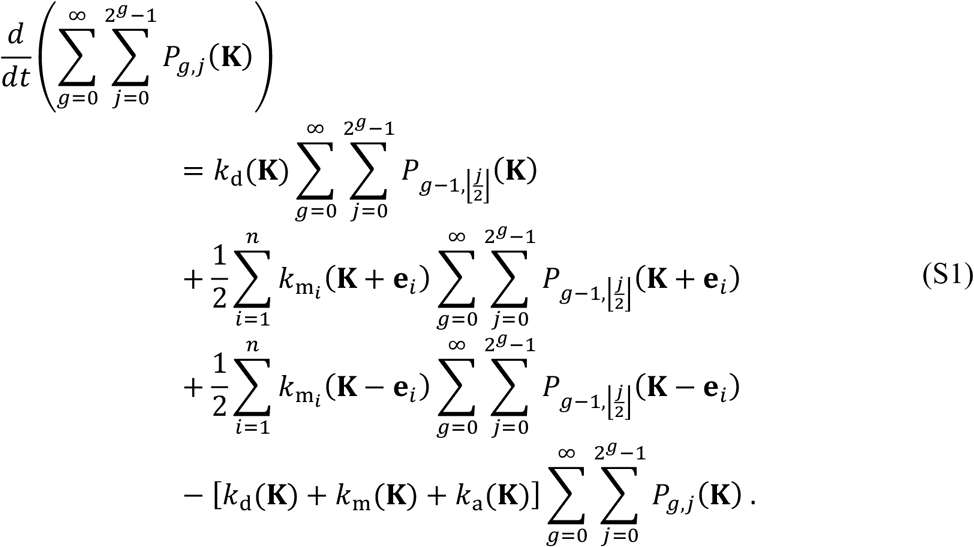

In the first term on the left-hand side (LHS) and last term on the right-hand side (RHS) we recognize the definition of the number of cells 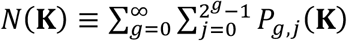. To calculate the expression 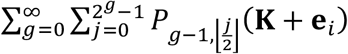 we first include the substitution *f* = *g* − 1,

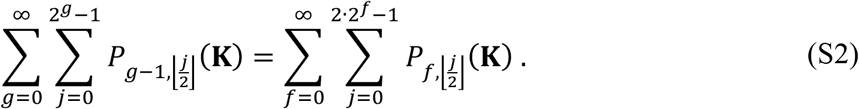

Note that summation starts with value *f* = 0 because negative generations do not exist, 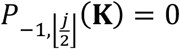. The floor function, 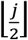, has the same value for two values of index *j*, and therefore we divide sum over index *j* into two sums, one for *j* = 2*a* and other for *j* = 2*b* + 1, each of them being equal to number of cells,

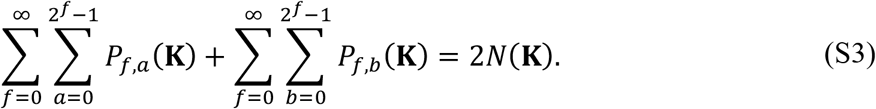

With this, Eq. (S1) reads,

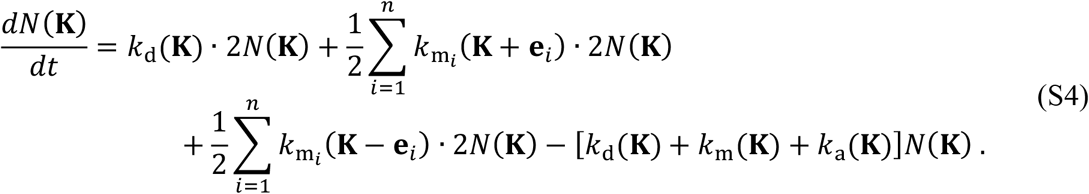

After rearranging Eq. (S4) we get the final form of the rate equation, Eq. (2).

### Calculations of rates of missegregation and apoptosis

In order to calculate rate of missegregation and apoptosis, that we use in our mean field approach, from corresponding probabilities, we integrate Eq. (2) and compare obtained number of cells with number of cells calculated from discrete cell divisions. In the case when initially all cells are diploids, number of cells with karyotypes **K**_2*n*_ ± **e**_***i***_ is negligible during first generation, and thus we ignore these contributions in Eq. (2) yielding

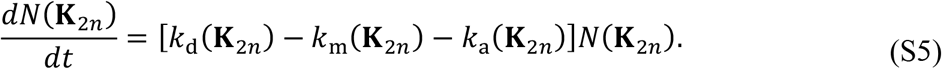

Because all terms in Eq. (S5) depend on the same karyotype, **K**_.2*n*_, we omit it from our notation in the rest of the derivation. By using expression *k*_d_ + *k*_m_ + *k*_a_ = *β* and the definition for rates of missegregation and apoptosis, 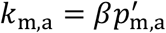, where 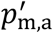 denotes the effective probabilities for missegregation and apoptosis, our equation reads,

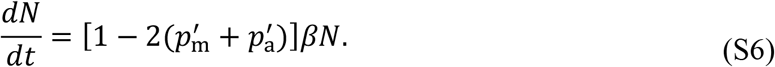

This equation has analytical solution, and number of cells after time *t*_0_ equals 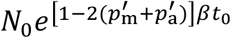. By taking into account expression *βt*_0_ = l*n* 2 we obtain,

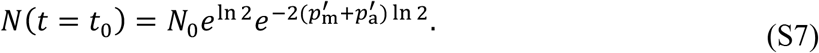

Finally, after linear approximation of the exponential function we get,

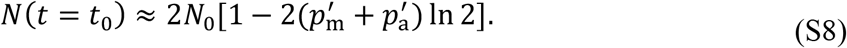

On the other hand, in our stochastic approach cell divisions occur after time *t*_D_, and number of cells is given by *N*_stoch._ = 2*N*_0_ [1 − (*p*_m_ + *p*_a_)]. Because number of cells should be equal by using both approaches, effective probabilities are given by 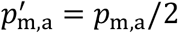, yielding the corresponding rates *k*_m,a_ = *βp*_m,a_ /2l*n*2.

### Macro-karyotype rate equation

The macro-karyotype is key tool in reducing the dimensionality of karyotype vector space, therefore we derive the rate equation for macro-karyotype evolution. Macro-karyotype is a vector **M** = (*x*_1_, …, *x*_*L*_), which components 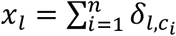 denote number of chromosomes with *l* copies, where *l* = 1, …, *L*. Index *L* denotes the maximal copy number of each chromosome. Note that all permutations of one set of components *c*_1_, …, *c*_*n*)_, belong to the same macro-karyotype.

In order to derive the rate equation for macro-karyotype we sum Eq. (2) over chromosome permutations,

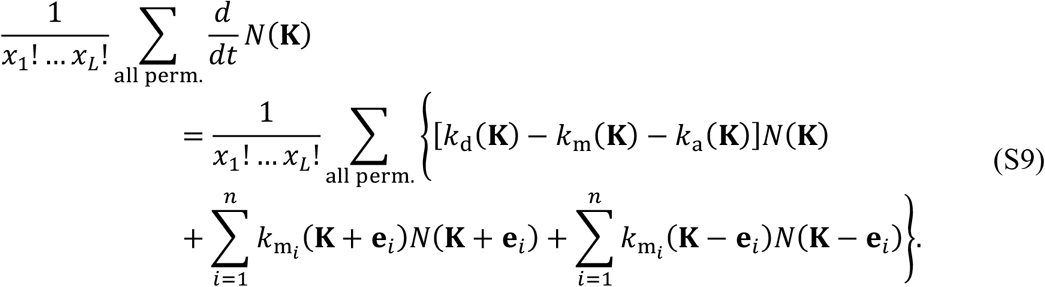

Here operator ∑_all perm._ denotes sum over all permutations of chromosome copy numbers *c*_1_, …, *c*_*n*)_ for a given karyotype **K**, whereas in the case of composed karyotypes, such as **K** + **e**_*i*_ = (*c*_1_, …, *c*_*i*_ + 1, …, *c*_*n*_), it denotes sum over all permutations of the composed karyotype.

It is useful to define number of cells that belong to the same macro-karyotype, Ñ(**M**) = (*x*_1_ ! … *x*_*L*_ !) ^−1^ ∑_all perm._ *N*(**K**). Equation (S9) simplifies dramatically in the case when the rates *k*_d,m,a_ (**K**) have the same value for all permutations of components of vector **K**, i.e. these rates have one value for the same macro-karyotype. We implement this constrain by introducing rates that are functions of macro-karyotype and have the same value as rates for corresponding karyotypes, 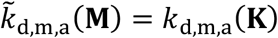, where **M** = **M**(**K**). In this case, the terms on the RHS simplify to (*x*_1_ ! … *x*_*L*_ !)^−1^ ∑_all perm.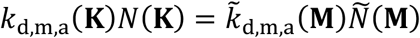_. The rate of missegregation for a given karyotype reads 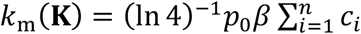. For a given macro-karyotype, this rate becomes 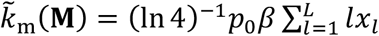. The sums stand for the total number of chromosomes, 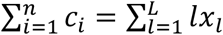.

In the case of the composed karyotype **K** + **e**_*i*_, that appears in the second term on the RHS, the number of cells of the associated macro-karyotype is 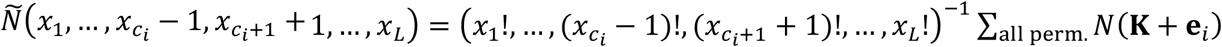. Similar applies for composed karyotype **K** – **e**_*i*_ in the third term. With this, our equation reads

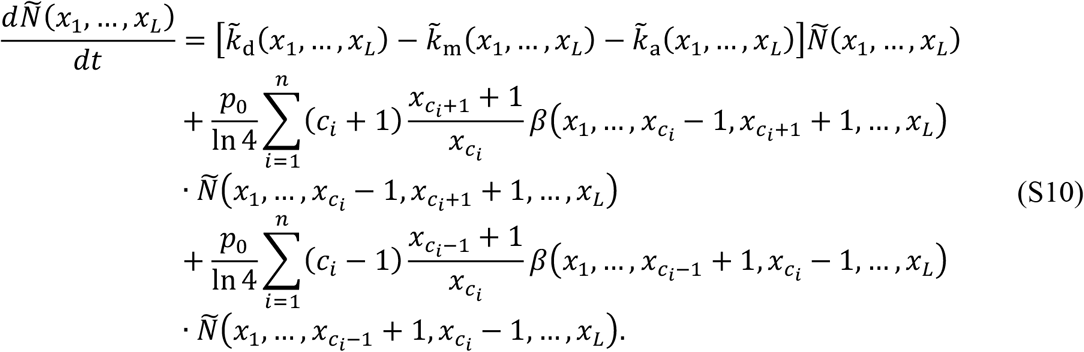

Here we used the definition of 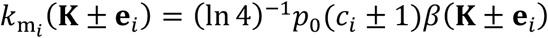. In order to group the elements with respect to the copy number we used the identity 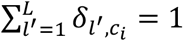, as well as the definition 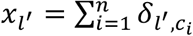, Eq. (S10) reads

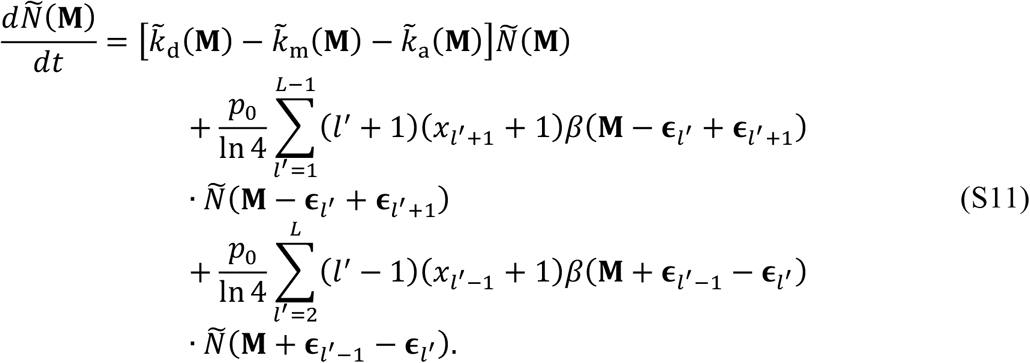

Note that in Eq. (S11) we write composed macro-karyotypes by using a unit vector, **ε**_e_′, which has the value 1 in the *l*′th coordinate and 0’s elsewhere. The upper summation limit in the second term is *L* − 1 in order to avoid copy numbers greater than maximal copy number *L*, and the lower summation limit in the third term is 2 because cells with copy number 0 are not viable. Lastly, we consider substitution *l* = *l*′ + 1 for the second and *l* = *l*′ − 1 for the third term with which we yield the final form of macro-karyotype rate equation,

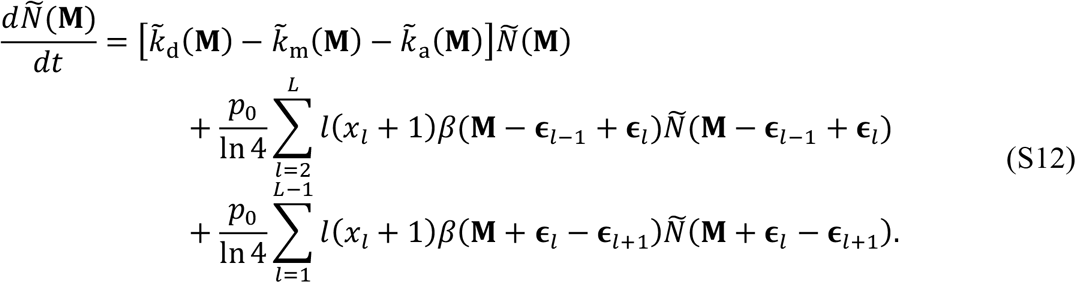

The final form of this rate equation is given in the main text, Eq. (3), where we recognize rate of chromosome loss 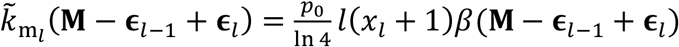 and similarly for the rates of chromosome gain.

For studying karyotypes with a specific chromosome, we generalize our approach by constructing an extended macro-karyotype **M**_ext_ (**K**) ≡ (*x*_1_, …, *x*_*L*_, *c*_15;_), where the last component, which corresponds to the specific chromosome, accounts for number of copies of chromosome 15 and rate equation for extended macro-karyotype yields,

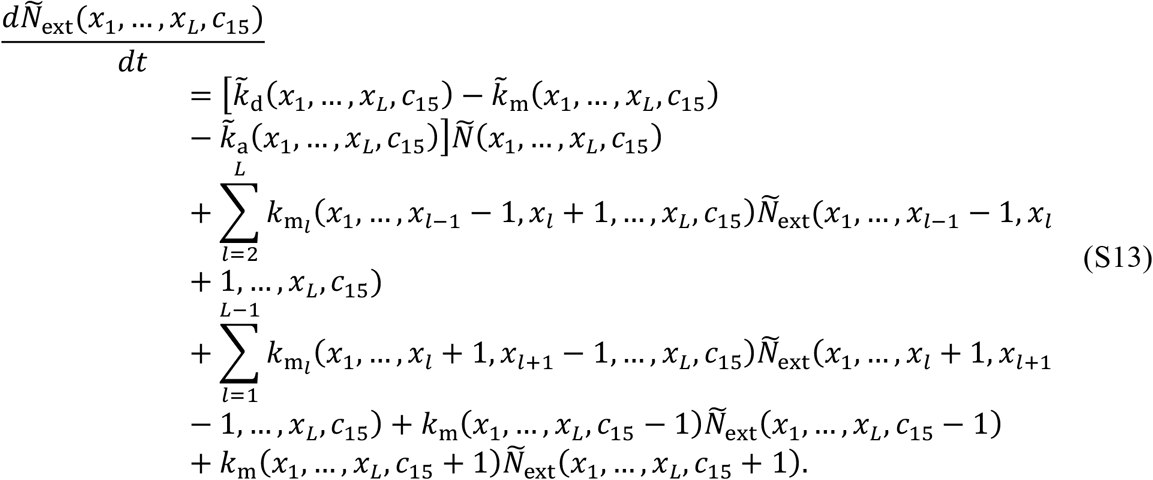

Where the rates 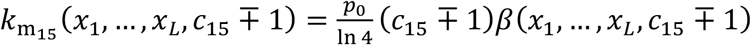 describe the missegregation of chromosome 15.

### Approximative form for chromosome gains

In order to estimate chromosome gains we calculate number of cells with three copies of *i*th chromosome, *N*(**K**_2*n*_ + **e**_*i*_), by using Eq. (2). Initially all cells are diploids and in the regime of a small missegregation probability and for small number of generations, where majority of cells have diploid karyotypes, *N*(**K**_2*n*_ + **e**_*i*_) ≪ *N*(**K**_2*n*_), Eq. (2) simplifies to,

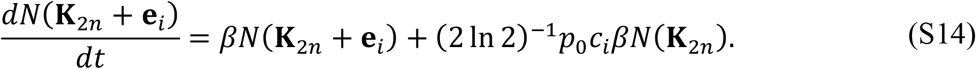

Number of cells *N*(**K**_2*n*_) in the second term on the RHS is obtained from rate equation *dN*(**K**_2*n*_)/*dt* = *βN*(**K**_2*n*_) and grows exponentially in time, *N*(**K**_2*n*_) = *N*_0_ *e* ^*β t*^, where *N*_0_ is initial number of cells. By solving Eq. (S14) we find the expression for number of cells with chromosome gains,

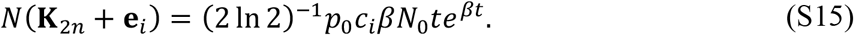

Generally, we define gain as the number of cells with gained chromosome with respect to total number of cells, ga*in* = *N*(**K**_2*n*_ + **e**_*i*_)/*N*(**K**_2*n*_) and in this case gain equals

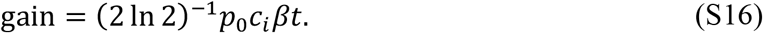

And finally, by considering that *β* = l*n* 2 /*t*_0_ and that *c*_*i*_ = 2 final form of equation for gain yields

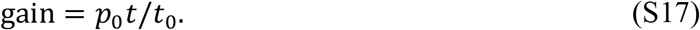

## REFERENCES

Araujo, A., Baum, B., and Bentley, P. (2013). The Role of Chromosome Missegregation in Cancer Development: A Theoretical Approach Using Agent-Based Modelling. PLOS ONE 8, e72206. https://doi.org/10.1371/journal.pone.0072206.

Bakhoum, S.F., Genovese, G., and Compton, D.A. (2009). Deviant Kinetochore Microtubule Dynamics Underlie Chromosomal Instability. Current Biology 19, 1937–1942. https://doi.org/10.1016/j.cub.2009.09.055.

Ben-David, U., and Amon, A. (2019). Context is everything: aneuploidy in cancer. Nature Reviews Genetics 21, 44–62. https://doi.org/10.1038/s41576-019-0171-x.

Ben-David, U., Arad, G., Weissbein, U., Mandefro, B., Maimon, A., Golan-Lev, T., Narwani, K., Clark, A.T., Andrews, P.W., Benvenisty, N., et al. (2014). Aneuploidy induces profound changes in gene expression, proliferation and tumorigenicity of human pluripotent stem cells. Nature Communications 5, 4825. https://doi.org/10.1038/ncomms5825.

Bolhaqueiro, A.C.F., Ponsioen, B., Bakker, B., Klaasen, S.J., Kucukkose, E., van Jaarsveld, R.H., Vivié, J., Verlaan-Klink, I., Hami, N., Spierings, D.C.J., et al. (2019). Ongoing chromosomal instability and karyotype evolution in human colorectal cancer organoids. Nature Genetics 51, 824–834. https://doi.org/10.1038/s41588-019-0399-6.

Bolton, H., Graham, S.J.L., van der Aa, N., Kumar, P., Theunis, K., Fernandez Gallardo, E., Voet, T., and Zernicka-Goetz, M. (2016). Mouse model of chromosome mosaicism reveals lineage-specific depletion of aneuploid cells and normal developmental potential. Nature Communications 7, 11165. https://doi.org/10.1038/ncomms11165.

Cimini, D. (2008). Merotelic kinetochore orientation, aneuploidy, and cancer. Biochimica et Biophysica Acta - Reviews on Cancer 1786, 32–40. https://doi.org/10.1016/j.bbcan.2008.05.003.

Cimini, D., Tanzarella, C., and Degrassi, F. (1999). Differences in malsegregation rates obtained by scoring ana-telophases or binucleate cells. Mutagenesis 14, 563–568. https://doi.org/10.1093/mutage/14.6.563.

Cimini, D., Howell, B., Maddox, P., Khodjakov, A., Degrassi, F., and Salmon, E.D. (2001). Merotelic Kinetochore Orientation Is a Major Mechanism of Aneuploidy in Mitotic Mammalian Tissue Cells 7. The Journal of Cell Biology 153, 517–528. https://doi.org/10.1083/jcb.153.3.517.

Desper, R., Difilippantonio, M.J., Ried, T., and Schäffer, A.A. (2005). A comprehensive continuous-time model for the appearance of CGH signal due to chromosomal missegregations during mitosis. Mathematical Biosciences 197, 67–87. https://doi.org/10.1016/j.mbs.2005.05.005.

Dewhurst, S.M., McGranahan, N., Burrell, R.A., Rowan, A.J., Grönroos, E., Endesfelder, D., Joshi, T., Mouradov, D., Gibbs, P., Ward, R.L., et al. (2014). Tolerance of whole-genome doubling propagates chromosomal instability and accelerates cancer genome evolution. Cancer Discovery 4, 175–185. https://doi.org/10.1158/2159-8290.CD-13-0285.

Drost, J., and Clevers, H. (2018). Organoids in cancer research. Nature Reviews Cancer 18, 407–418. https://doi.org/10.1038/s41568-018-0007-6.

Duijf, P.H.G., and Benezra, R. (2013). The cancer biology of whole-chromosome instability. Oncogene 32, 4727–4736. https://doi.org/10.1038/onc.2012.616.

Elizalde, S., Laughney, A.M., and Bakhoum, S.F. (2018). A Markov chain for numerical chromosomal instability in clonally expanding populations. PLoS Computational Biology 14, e1006447. https://doi.org/10.1371/journal.pcbi.1006447.

Ganem, N.J., Godinho, S.A., and Pellman, D. (2009). A mechanism linking extra centrosomes to chromosomal instability. Nature 460, 278–282. https://doi.org/10.1038/nature08136.

Gordon, D.J., Resio, B., and Pellman, D. (2012). Causes and consequences of aneuploidy in cancer. Nature Reviews Genetics 13, 189–203. https://doi.org/10.1038/nrg3123.

Gusev, Y., Kagansky, V., and Dooley, W.C. (2000). A Stochastic Model of Chromosome Segregation Errors with Reference to Cancer Cells. Mathematical and Computer Modelling 32, 97–111. https://doi.org/10.1016/S0895-7177(00)00122-9.

Gusev, Y., Kagansky+, V., and Dooley, W.C. (2001). Long-Term Dynamics of Chromosomal Instability in Cancer: A Transition Probability Model. Mathematical and Computer Modelling 33, 1253–1273. https://doi.org/10.1016/S0895-7177(00)00313-7.

Heyde, A., Rohde, D., McAlpine, C.S., Zhang, S., Hoyer, F.F., Gerold, J.M., Cheek, D., Iwamoto, Y., Schloss, M.J., Vandoorne, K., et al. (2021). Increased stem cell proliferation in atherosclerosis accelerates clonal hematopoiesis. Cell 184, 1348-1361.e22. https://doi.org/10.1016/j.cell.2021.01.049.

Hintzen, D.C., Soto, M., Schubert, M., Bakker, B., Spierings, D.C., Szuhai, K., Lansdorp, P.M., Foijer, F., Medema, R.H., and Raaijmakers, J.A. (2021). Monosomies, trisomies and segmental aneuploidies differentially affect chromosomal stability. BioRxiv https://doi.org/10.1101/2021.08.31.458318.

Hwang, S., Cavaliere, P., Li, R., Zhu, L.J., Dephoure, N., and Torres, E.M. (2021). Consequences of aneuploidy in human fibroblasts with trisomy 21. Proc Natl Acad Sci U S A 118, e2014723118. https://doi.org/10.1073/pnas.2014723118.

van Jaarsveld, R.H., and Kops, G.J.P.L. (2016). Difference Makers: Chromosomal Instability versus Aneuploidy in Cancer. Trends in Cancer 2, 561–571. https://doi.org/10.1016/j.trecan.2016.09.003.

Jelenic, I., Selmecki, A., Laan, L., and Pavin, N. (2018). Spindle Dynamics Model Explains Chromosome Loss Rates in Yeast Polyploid Cells. Frontiers in Genetics 9, 296. https://doi.org/10.3389/fgene.2018.00296.

Laughney, A.M., Elizalde, S., Genovese, G., and Bakhoum, S.F. (2015). Dynamics of Tumor Heterogeneity Derived from Clonal Karyotypic Evolution. Cell Reports 12, 809–820. https://doi.org/10.1016/j.celrep.2015.06.065.

Lengauer, C., Kinzler, K., and Vogelstein, B. (1997). Genetic instability in colorectal cancers. Nature 386, 623–627. https://doi.org/https://doi.org/10.1038/386623a0.

Liu, Y., Sethi, N.S., Hinoue, T., Schneider, B.G., Cherniack, A.D., Sanchez-Vega, F., Seoane, J.A., Farshidfar, F., Bowlby, R., Islam, M., et al. (2018). Comparative Molecular Analysis of Gastrointestinal Adenocarcinomas. Cancer Cell 33, 721-735.e8. https://doi.org/10.1016/j.ccell.2018.03.010.

Lynch, A.R., Arp, N.L., Zhou, A.S., Weaver, B.A., Burkard, M.E., and Burkard, M. (2021). Quantifying chromosomal instability from intratumoral karyotype diversity using agent-based modeling and Bayesian inference. BioRxiv.

Lynch, A.R., Arp, N.L., Zhou, A.S., Weaver, B.A., and Burkard, M.E. (2022). Quantifying chromosomal instability from intratumoral karyotype diversity using agent-based modeling and Bayesian inference. Elife 11, e69799. https://doi.org/10.7554/eLife.69799.

Narkar, A., Johnson, B.A., Bharne, P., Zhu, J., Padmanaban, V., Biswas, D., Fraser, A., Iglesias, P.A., Ewald, A.J., and Li, R. (2021). On the role of p53 in the cellular response to aneuploidy. Cell Reports 34, 108892. https://doi.org/10.1016/j.celrep.2021.108892.

Nicholson, J.M., and Cimini, D. (2013). Cancer karyotypes: Survival of the fittest. Frontiers in Oncology 3, 148. https://doi.org/10.3389/fonc.2013.00148.

Nicholson, J.M., Macedo, J.C., Mattingly, A.J., Wangsa, D., Camps, J., Lima, V., Gomes, A.M., Ried, T., Logarinho, E., and Cimini, D. (2015a). Chromosome mis-segregation and cytokinesis failure in trisomic human cells. ELIFE 4, e05068. https://doi.org/10.7554/eLife.05068.

Nicholson, J.M., Macedo, J.C., Mattingly, A.J., Wangsa, D., Camps, J., Lima, V., Gomes, A.M., Dória, S., Ried, T., Logarinho, E., et al. (2015b). Chromosome mis-segregation and cytokinesis failure in trisomic human cells. Elife 4, e05068. https://doi.org/10.7554/eLife.05068.

Petrie, H.T., and Zúñiga-Pflücker, J.C. (2007). Zoned out: functional mapping of stromal signaling microenvironments in the thymus. Annual Review of Immunology 25, 649–679. https://doi.org/10.1146/annurev.immunol.23.021704.115715.

Porritt, H.E., Gordon, K., and Petrie, H.T. (2003). Kinetics of steady-state differentiation and mapping of intrathymic-signaling environments by stem cell transplantation in nonirradiated mice. Journal of Experimental Medicine 198, 957–962. https://doi.org/10.1084/jem.20030837.

Prasad, K., Bloomfield, M., Levi, H., Keuper, K., Bernhard, S. v, Baudoin, N.C., Leor, G., Eliezer, Y., Giam, M., Wong, C.K., et al. (2022). Whole-Genome Duplication Shapes the Aneuploidy Landscape of Human Cancers. Cancer Res 82, 1736–1752. https://doi.org/10.1158/0008-5472.CAN-21-2065/681796/AM/WHOLE-GENOME-DUPLICATION-SHAPES-THE-ANEUPLOIDY.

Rohban, S., and Campaner, S. (2015). Myc induced replicative stress response: How to cope with it and exploit it. Biochimica et Biophysica Acta 1849, 517–524. https://doi.org/10.1016/j.bbagrm.2014.04.008.

Santaguida, S., Vasile, E., White, E., and Amon, A. (2015). Aneuploidy-induced cellular stresses limit autophagic degradation. Genes and Development 29, 2010–2021. https://doi.org/10.1101/gad.269118.115.

Sheppard, O., Wiseman, F.K., Ruparelia, A., Tybulewicz, V.L.J., and Fisher, E.M.C. (2012). Mouse Models of Aneuploidy. Scientific World Journal 2012, 214078. https://doi.org/10.1100/2012/214078.

Shi, Q., and King, R.W. (2005). Chromosome nondisjunction yields tetraploid rather than aneuploid cells in human cell lines. Nature 437, 1038–1042. https://doi.org/10.1038/nature03958.

Shoshani, O., Bakker, B., de Haan, L., Tijhuis, A.E., Wang, Y., Kim, D.H., Maldonado, M., Demarest, M.A., Artates, J., Zhengyu, O., et al. (2021). Transient genomic instability drives tumorigenesis through accelerated clonal evolution. Genes & Development 35, 1093–1109. https://doi.org/10.1101/gad.348319.121.

Storchova, Z., and Kuffer, C. (2008). The consequences of tetraploidy and aneuploidy. Journal of Cell Science 121, 3859–3866. https://doi.org/10.1242/jcs.039537.

Thompson, S.L., and Compton, D.A. (2008). Examining the link between chromosomal instability and aneuploidy in human cells. Journal of Cell Biology 180, 665–672. https://doi.org/10.1083/jcb.200712029.

Thompson, S.L., and Compton, D.A. (2011a). Chromosome missegregation in human cells arises through specific types of kinetochore-microtubule attachment errors. Proc Natl Acad Sci U S A 108, 17974–17978. https://doi.org/10.1073/pnas.1109720108.

Thompson, S.L., and Compton, D.A. (2011b). Chromosome missegregation in human cells arises through specific types of kinetochore-microtubule attachment errors. Proc Natl Acad Sci U S A 108, 17974–17978. https://doi.org/10.1073/pnas.1109720108.

Tolic, I.M., and Pavin, N. (2021). Mitotic spindle: Lessons from theoretical modeling. Molecular Biology of the Cell 32, 218–222. https://doi.org/10.1091/mbc.E20-05-0335.

Trakala, M., Aggarwal, M., Sniffen, C., Zasadil, L., Carroll, A., Ma, D., Su, X.A., Wangsa, D., Meyer, A., Sieben, C.J., et al. (2021). Clonal selection of stable aneuploidies in progenitor cells drives high-prevalence tumorigenesis. Genes and Development 35, 1079– 1092. https://doi.org/10.1101/gad.348341.121.

Weaver, B.A., and Cleveland, D.W. (2006). Does aneuploidy cause cancer? Current Opinion in Cell Biology 18, 658–667. https://doi.org/10.1038/nrg3123.

Williams, B.R., Prabhu, V.R., Hunter, K.E., Glazier, C.M., Whittaker, C.A., Housman, D.E., and Amon, A. (2008). Aneuploidy Affects Proliferation and Spontaneous Immortalization in Mammalian Cells. Science (1979) 322, 703–708. https://doi.org/10.1126/science.1160058.

